# A simple pyrocosm for studying soil microbial response to fire reveals a rapid, massive response by *Pyronema* species

**DOI:** 10.1101/763169

**Authors:** Thomas D. Bruns, Judy A. Chung, Akiko A. Carver, Sydney I. Glassman

## Abstract

We have designed a simple, inexpensive system for the studying the response of soil microbes to fire. This system allows one to create post-fire environments in soil in reproducible and realistic ways. Using it we show that the peak soil temperature achieved at a given depth occurs hours after the fire is out, lingers near peak temperature for a significant time, and is accurately predicted by the log of soil depth and the mass charcoal burned. Flash fuels that left no large coals were found to have a negligible soil heating effect. Coupling this system with Illumina MiSeq sequencing of the control and post-fire soil we show that we can stimulate a rapid, massive response by *Pyronema*, a well-known genus of postfire fungus, from uninoculated forest soil within two weeks of a test fire. This specific stimulation occurs in a background of many other fungal taxa that do not change significantly with the fire, although there is an overall reduction in richness and evenness. Extrapolating from the physical relationships we predict soil heating effects in wild fires are likely to be very patchy across the forest floor but the width of a survivable “goldilocks zone” will stay relatively constant across a range of fuel loads. We further predict that a necromass zone above it, which represents an open niche for pyrophilous microbes, increases in size rapidly with addition of fuel, and then remains nearly constant over a broad range of fuel loads. The simplicity of this experimental system, coupled with the availability of a set of sequenced, assembled and annotated genomes of pyrophilous fungi, offers a powerful tool for dissecting the ecology of post-fire microbial communities.

## Introduction

Fire is a natural part of many ecosystems, and organisms in systems with predictable fire regimes are often well adapted to survive or recolonize rapidly after fire. Plant adaptions are particularly well known with either the ability to survive fire through thickened bark, sertotinous pine cones, vegetative resprouting, or other traits [1]. Soil microbes also must survive or recolonize after fires, but much less is known about how they achieve this or what their roles are in the post-fire environment. Pressler et al. [2] conducted a meta-analysis on belowground effects of fire and reported that negative effects are commonly found on microbes including reductions in biomass, abundance, richness and evenness across taxonomic groups, and these effects were coupled with decade-long recovery times. Fungi in particular showed large reductions and slow recovery [2]. Dove and Hart [3] found similar effects in their meta-analysis of only fungal communities, and both studies showed that the largest effects were associated with forest biomes. Post-fire changes in fungal communities were thought to be caused by both the direct killing effects of fire on the organisms, and by indirect effects on habitat, such as the loss of the organic layers and the reduction in root biomass of mycorrhizal hosts.

Almost all studies considered in these meta-analyses focused on the reduction in post-fire microbial communities, but there is also evidence of microbes that respond positively to fire. In particular there is a set of “**pyrophilous**” fungi that fruit only in burnt habitats and are abundant in the first weeks or months following fire. These saprophytic fungi have been known for over a hundred years [4], and the pattern of their fruiting in post-fire settings suggests rapid successional sequences [5, 6]. Some of these fungi have heat-stimulated spores [7], or spores that can also be stimulated by post-fire chemicals [8].

Nevertheless, the rapid, post-fire, fungal succession has not been confirmed at the mycelial level with modern, molecular methods, nor is it known what these fungi are doing in post-fire environment.

Similar to the saprobic communities, most ectomycorrhizal fungi are reduced, but a few thrive in the post-fire environment. In stand-replacing fires, composition of ectomycorrhizal communities shifts from the resident community that dominated the roots on mature trees to a community that survived in the “spore bank” and colonizes establishing seedlings [9, 10]. Some component species in the spore bank have heat-resistant spores [11] and exhibit an increase in abundance in post-fire or experimentally heated soils [10, 12, 13]. This adds credence to the idea that some microbes are adapted to fires.

The direct heating effects of fire and production of fire-specific soil chemistry are likely to interact with post-fire microbial communities in multiple ways. Fire heating of soils is reasonably well understood from a variety of models [14–16]. These show that the heat capacity and water content of soil produce depth-stratified temperatures. The effects of fire on soil chemistry are also understood in general ways, and are known to be correlated with heating and depth. For example, extreme surface temperatures can completely combust much of the ligno-cellulosic biomass into CO_2_, while producing a cation-rich, high pH ash. Substantial amounts of biomass are also transformed into partially burned “pyrolyzed” carbon sources [17]. Temperatures from 480-220 °C convert biomass into a mixture of partially pyrolyzed organic compounds, while volatilized waxes and lipids typically condense at lower temperatures in the soil below. This process results in the hydrophobic soils that commonly occur in forest fires and can lead to heavy erosion by channelizing runoff [18–20]. This erosion reduces site productivity and is one of the greatest concerns for post-fire recovery. Hydrophobic post-fire soils can last years or disappear in weeks or months, but the biotic or abiotic processes that determine their degradation are unknown. At lower depths, where peak temperature is below 200 °C, little pyrolysis occurs, but the high temperatures still kill most life. This creates a necromass zone, where dead organisms leak out easily-mineralizable forms of carbon, creating a carbon subsidy for any microbes that can survive or rapidly recolonize this zone.

Experimental evidence of the effects of fire on microbial communities has been largely limited to sampling studies of wild-fires and prescribed burns (see studies included in [2, 3]). Although some generalities have been learned from these approaches, they have not been useful for connecting the detailed understanding of soil heating and chemistry to structure of post-fire microbial communities. In addition post-fire successional studies have generally been space-for-time comparisons with limited replication, and sampling was usually not conducted until at least 1 year and typically several years post fire [2, 3]. Thus, initial changes in fungal communities were missed, or if in studies in which early sampling occurred, they were conducted prior to molecular identification methods and relied on fruiting [6].

To create a more experimental approach we have developed a “**pyrocosm**” system that allows soil samples to be heated by fire while the temperatures are monitored. These samples can then be incubated as intact units, or targeted temperature/depth zones can be removed and inoculated with test organisms *in vitro*. Here we report the thermal characteristics of this system, and show that they can be used to stimulate native pyrophilous fungi in test soils.

## Materials and Methods

### Pyrocosm design

Our pyrocosms consisted of 1 gallon galvanized steel buckets, filled to a depth of 16cm with test soil or sand, wired with K-type thermocouples at various soil depths, and buried to soil level at the Oxford Tract at the University of California Berkeley. Total volume of soil in the buckets was 7 liters. Ten to 13 drill holes (0.6 cm diameter) above the soil surface were added to increase aeration for the fire. Additional smaller diameter drill holes in the side of bucket were used to place thermocouples into selected depths. A small fire with weighed fuels consisting of pine needle litter, paper and charcoal briquettes was then burned on the top of it (Fig 1) and the test soil was watered the next day with to initiate microbial activity after it has completely cooled. Additional notes on assembling these pyrocosms are given in the supplemental material (File S1)

**Fig 1.**
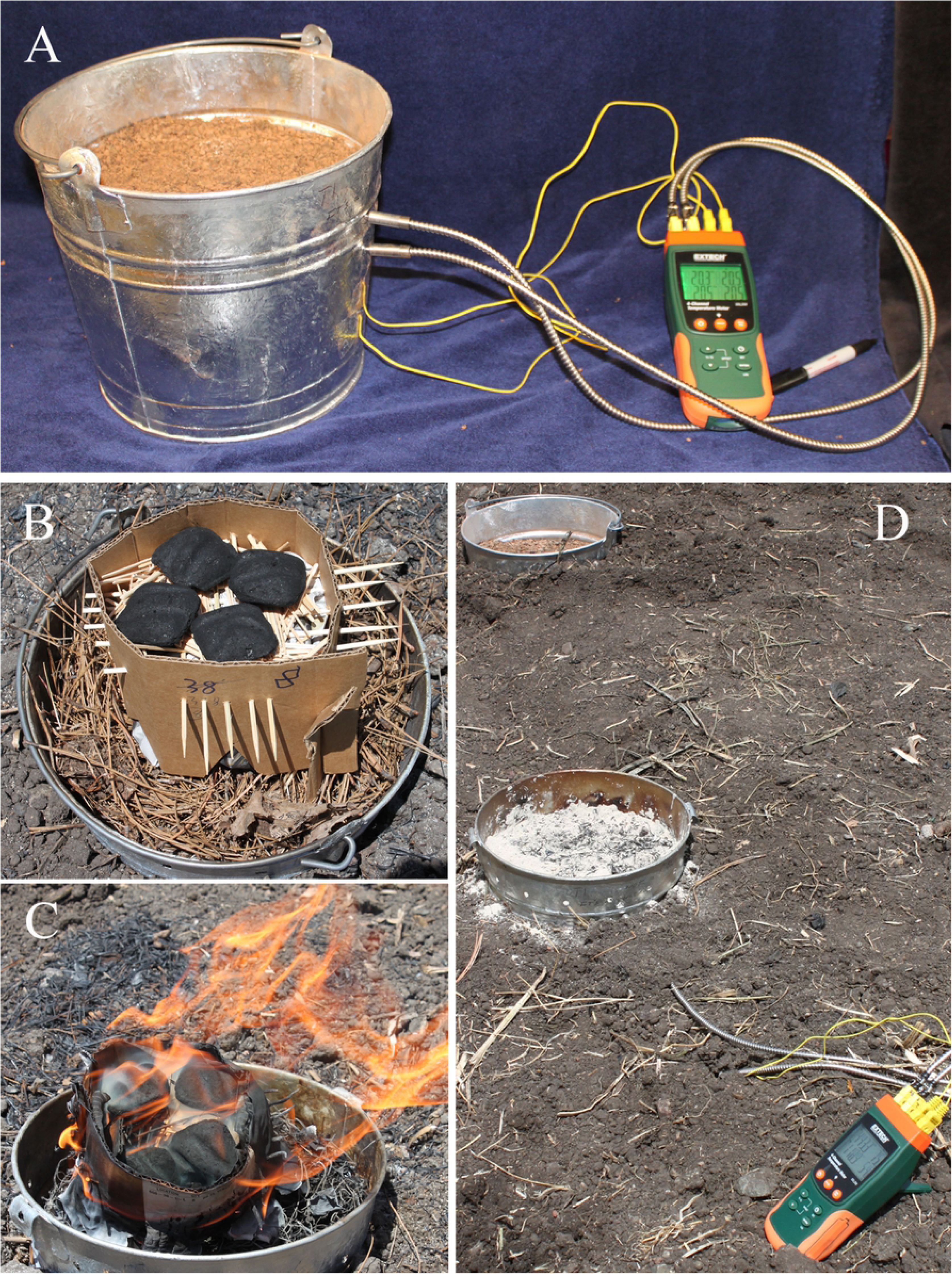
Pyrocosm construction. A) Pyrocosm in the lab filled with forest soil and wired with thermocouples; air holes around rim had not yet been drilled but are seen in remaining images; B) weighted cardboard, tooth picks and newspaper constitute “flash fuels” used to ignite charcoals; C) early ignition phase; D) burned pyrocosm and unburned control in background.

The two pyrocosms used to study post-fire fungal response used forest soil. Unburned test soil for these was collected from a *Pinus ponderosa* forest in Stanislaus National forest, CA, USA, 37.8140 - 120.0689, just outside the perimeter of the 2013 Rim Fire. The top 5 cm of soil, and the bottom 5-15 cm were collected separately. Burned test soil was collected from a nearby, fire-killed, *Pinus ponderosa* forest within the perimeter of the Rim fire in Stanislaus National forest, 37.8442 - 119.9402, located adjacent to previously studied plots [10], and was not depth stratified. All soils were collected in Spring 2015. Litter and F layers were collected and combined from the same unburned site. The soils were sieved through a 2mm soil screen to homogenize them and remove rocks, roots and large aggregates, and then were air-dried in large, closed paper bags for 3 weeks in a fume hood to a moisture content of between 3.8% (bottom 5-15 cm) and 4.8% (top 5 cm), and then stored in sealed plastic bags at 5° C until needed. When assembling soil pyrocosms, bottom soil was used for the lower 5-16 cm and top soil was used for the upper 5 cm. In the second soil pyrocosm, burned forest soil was mixed with the bottom unburned soil in a 1:9 ratio with intent of adding inoculum of post-fire fungi to the experiment.

Course sand (∼2mm grain) was used instead of soil in the series of 14 experiments that tested the relationship between fuel and soil temperatures. Sand was selected because it was not altered by the fire, and therefore pyrocosm experiments could be repeated without reassembling the units between fires. Instead, the ash and charcoal formed were simply removed by hand and with the assistance of a small fan, and then a new fuel load and heating experiment were tested. Controls for these temperature experiments consisted of an unburned, sand pyrocosm that was located 1m from the burned sand pyrocosms. This gave us a reading of diurnal temperature changes unrelated to the fires.

Thermocouples were K-type, with an accuracy of 1°C. In the first soil pyrocosms, four thermocouples were installed through drilled holes in the side of the buckets at 0.5, 3.5, 6.5, and 9.5 cm from the surface (Fig 1a). These were inserted as the soil was added such that the tip of each thermocouple was located near the center of the bucket at the prescribed depths (see file S1 for detailed notes). The top two thermocouples were insulated K-type thermocouples that could withstand temperatures up to 1038°C (custom made by OMEGA.com, see Fig 1A). The lower two were typical, plastic insulated wire thermocouples with temperature maximums of 200°C (Fig 1a). In later experiments, the number and arrangement of thermocouples per pyrocosm were modified as needed for specific uses. For experiments designed to determine the effect of fuel load on peak temperatures only two thermocouples were used and placed at 10.5 and 15.8 cm from the surface. These were the plastic insulated types since temperatures were more moderate at these depths and the thinner wires were more accurately placed. Temperatures were recorded using a data logger (Extech Instruments SDL200) at 10 sec or 30 sec intervals. The 30 sec intervals turned out to be more than sufficient because temperatures change fairly gradually at the depths measured.

### Fuels, ignition and fire

Fuel in all experiments was weighed prior to ignition. Dried, weighed, forest litter and F-layer were placed at the soil surface and gently tamped down to achieve an approximate depth of approximately 4 cm, which was similar to that at the site where the soil and litter were collected. Litter was re-dried the night before use to ensure that it burned readily and was consumed completely. A weighed, but variable number of charcoal briquettes, obtained from a local grocery store, were placed on top of the litter.

The fire for the two soil pyrocosms contained approximately 300 g of litter and 1000 g of charcoal. The charcoal was piled in the middle of bucket, and was spread evenly after it was well lit. The temperatures at the four depths were monitored and the charcoal was then carefully removed with a small trowel when soil reached a target temperature. The removed charcoal was extinguished rapidly by dropping it in water, and was later dried and weighed and used to determine the amount that had been consumed. The temperature was monitored for six more hours and pyrocosms 1 and 2 ultimately reached temperatures of 146°C and 171°C, respectively, at a depth of 6.5 cm.

In all sand experiments the amount of charcoal was reduced to 1 to 15 briquettes (∼25 to 380 gm) and was ignited by perching it on top of fine wood kindling (tooth picks), held in a 28-38 g cardboard ring that was filled with a single sheet of newspaper (Fig 1b). After the cardboard ring burned, and the ignited charcoal was moved toward the center, spread evenly, and allowed to burn completely.

### Fungal incubation, sampling, and controls

The two pyrocosms filled with forest soil were used to study fungal response to fire. The day following an experimental fire, after the soil temperatures had returned to normal, water was added to the pyrocosms in the form of weighed, crushed ice. This method for adding the water was meant to mimic snow, which is a typical way late fall precipitation occurs at mid elevation in the Sierra Nevada Mountains where the soil was collected. The slow melting also allowed the water to gently and uniformly percolate into the soil, even if the soil had become slightly hydrophobic from the fire. Pyrocosm 1 received 1000 g of ice on day 1, and 500 g on day 2, Pyrocosm 2 received 1500 g of ice on day one and 1,500g on day 3. A control pyrocosm, filled with the same forest soil, setup in the same way and buried a meter from the test pyrocosm 1, was left unburned, but watered and incubated in the same way as the first pyrocosm (Fig 1D). After burning and watering pyrocosms were covered with foil to prevent additional precipitation, and to limit aerial dispersal, and were incubated *in situ*.

Soil within the pyrocosms was sampled with a 2cm diameter soil corer at one week and two weeks postfire for both test pyrocosms and in the control pyrocosm. A four week sample was also taken from the second experimental pyrocosm. Soil cores were vertically divided into four approximately even zones, and DNA was extracted from each via MoBio DNAeasy Power Soil kit (Qiagen, Carlsbad, CA, USA).

### Construction of Illumina libraries, and processing of Illumina data

We amplified ITS1 spacer, which is part of the Internal Transcribed Spacer region, the universal DNA barcode for fungi [21], with the ITS1F-ITS2 primer pair using Illumina sequencing primers designed by Smith and Peay [22] and prepared libraries for Illumina MiSeq PE 2 x 250 sequencing as previously described [10]. Sequencing was performed at the Genome Center at the University of California, Davis, CA, USA. Bioinformatics was performed with UPARSE [23] usearch v7 and QIIME 1.8 [24] with the same methods as previously published [10]. All analyses are based on 97% sequence similarity for operational taxonomic units (**OTUs**) and taxonomy was assigned with the UNITE fungi database [25] accessed on 30 Dec 2014. Samples for a related study were run at the same time and OTUs were processed for both simultaneously to enable later comparison. As a result OTU identifying numbers reported here are greater than the 887 total found in this study. Representative sequences for all OTU that represented 1% or more of the read abundance in either of the pyrocosms or the control were BLASTed [26] individually against the NCBI database, and the results were examined to improve the automated sequence-based identifications. Sequences are available at the NCBI Sequence Read Archive (PRJNA559408), and representative sequences from each of the OTUs are deposited in NCBI XXXXXXX.

A mock community composed of equal quantities of DNA extractions of *Coprinellus sp. Pholiota sp. Cyathus stercoreus, Penicillium citreonigrum*, *Aureobasidium pullulans*, *Rhodotorula mucilaginosa*, *Cladosporium sp*., *Suillus sp*., *Peyronellaea glomerata*, and *Peyronellaea glomerata* was included to control for OTU processing, and a no DNA control was included to assay for spurious contamination and index jumping [27].

### Statistical analyses

Plotting of temperature changes over time was initially done with Microsoft Excel, and then repeated in R along with all modeling and correlation analyses [28]. The response of individual OTUs was investigated by examining their changes in sequence abundance with simple metrics in a excel spreadsheet (File S2). To determine percent sequence abundance for a given taxon at a particular time point, sequences for all depths at a given time point and experiment were summed, divided by the sum of all sequences of all taxa for that same time point of the experiment, and multiplied by 100. Ideally each time point of an experiment should have had 4 depth samples but four of the individual depth samples out of 28 did not yield sequences and were dropped from the further consideration. The dropped samples were: Pyrocosm 1, week 1, depth 3; Pyrocosm 2, week1, depths 1 and 2; and Pyrocosm 2 week 2, depth 4.

To determine overall abundance ranks of OTUs, the sequences across all depths in weeks one and two were separately summed for each OTU for each of the two pyrocosms and the control, and divided by total sequences of all OTUs for the experimental unit (pyrocosm 1, 2 or control). The spreadsheet was then sorted by sequence abundance, first for the control and then for each of the two experimental pyrocosms. Ranks were assigned in descending order within each experimental unit with the most abundant OTU as 1.

OTU increases or decreases relative to fire were investigated by subtracting percent sequence abundance for each OTU from each pyrocosm from the percent sequence abundance in the control. The OTUs were then sorted by this metric to identify those that showed the largest changes.

## Results

### Soil heating characteristics in the pyrocosms

The heating experiments showed the following about the peak temperature that is reached at a given depth: 1) The reproducibility of the temperature profiles is excellent when fuel levels are constant (Fig 2a). 2) Peak temperatures at depth are reached hours after the fire has gone out and they linger near the peak for 40 minutes or more (Fig 2a). 3) Peak soil temperatures decrease by the log of the depth (Fig 2b) and can be predicted by the mass of fuel (Fig 2c). 4). There are edge effects that result in progressively cooler temperatures away from the center (Fig 2d).

**Fig. 2.**
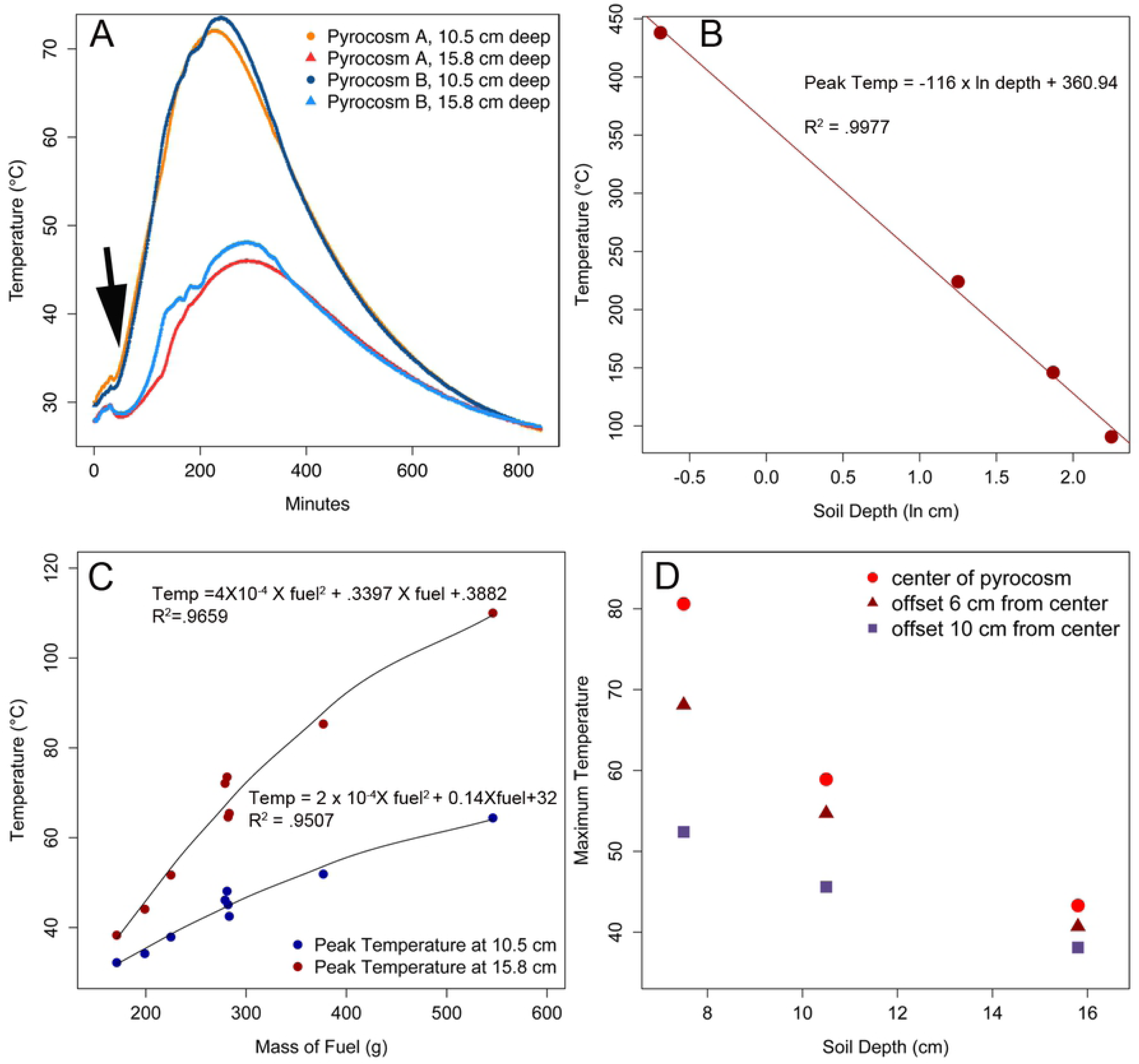
Thermal Characteristics of Pyrocosms. A) temperature profiles from two depths of two replicate pyrocosms. Traces plot half-minute intervals over 800 minutes. Note that that the temperature profiles are quite repeatable, and that temperature continues to rise after the fire is out (arrow); B) peak temperatures are predicted by the ln (or log10, not shown) of the depth, and C) by the mass of fuel; D) peak temperature at three depths (7.5, 10.5, 15.8 cm) and three locations offset from center within a depth. Note that the center is hottest and the edge of the unit is cooler.

A regression between peak temperatures and log of depth was linear with a R*^2^* of 0.9977 (Fig 2b). The predictability of peak temperatures and total fuel is shown as a polynomial model with R^2^ values of 0.9682 (10.5 cm depth), and 0.9507 (15.8 cm depth), but a simpler logarithmic model works nearly as well with R^2^ values of 0.9659 and 0.94779m respectively.

Flash fuels, those that leave little or no coals such as pine needles, small wooden sticks, cardboard, and newspaper, had almost undetectable heating effects on the soil at depth. Burning of 172 g of flash fuels (i.e., just litter, F-layer, paper, cardboard) caused a 2°C rise in temperature at 10.5 cm below the surface, relative to the unburned control. This contrasts with approximately an 8° C rise at the same depth by a single 25.5 g charcoal. However, the flash fuels did cause a more rapid rise in temperature than that caused by solar heating in the control (Fig S1), and using regression from the measured points the temperature 1 mm from the surface peaked at 108° C, and soil temperature was predicted to be heated to 70° C to a depth at a 1.26 cm.

There is a significant edge effect on soil heating in the pyrocosms, and it is most pronounced at the shallower depths (Fig. 2d). For example, at 7.5 cm the peak temperature varied from 91 to 61 °C, depending on whether the temperature was measured at the center or near the edge of the pyrocosm. Near the bottom of the unit at 15.8 cm, the temperature from center to edge only varied 5 ° from 48 to 43 ° C, which is a lesser absolute and relative difference than that recorded at 7.5 cm for the surface.

Spreading out the coals more did not greatly reduce this edge effect (data not shown). The edge effects do not affect the results presented above with respect the relationships between depth and mass of charcoal on peak temperature, or the temperature-time profiles except to limit their accuracy to the center of the units. However, these edge effect do mean that peak temperature isoclines would be curved upward on the edges.

### Fungal response to the pyrocosms

Table 1 summarizes the read depth, number of OTUs, and ranked abundance of the all OTUs in experimental and control samples that had sequence abundance of 1% or greater in control or pyrocosms. File S2 gives the complete OTU table in spreadsheet form with summary data included. Four of the 20 experimental samples yielded 58 or fewer sequences and were dropped. All other experimental samples yielded more than 6000 sequences and were retained. Because the dropped samples made the depth sampling difficult to compare across time, data from depth samples for single time points within an experimental unit were combined to give more robust measures of fungal communities within a time point for table 1.

**Table 1.** Summary of samples, read depth, and OTUs

|  | Unburned Controls | Pyrocasm 1 Weeks 1 & 2 | Pyrocasm 2 Weeks 1 & 2 | Pyrocasm 2 week 3 | Mock control | No DNA control |
| --- | --- | --- | --- | --- | --- | --- |
| # samples retained (and lost) | 8 | 7(1) | 6 (2) | 4 | 2 | 4 |
| Ave sequence depth/sample | 54427 | 67308 | 25247 | 8495 | 21087 | 408 |
| Range of sequence depth | 9639-71080 | 37014-109855 | 6607 - 68272 | 3950-15637 | 16530-25643 | 90-1044 |
| # OTUs >= 0.01% sequence | 406 | 158 | 155 | 110 | 24 | 20 |
| # OTUs with read depth > 5 seq | 605 | 296 | 191 | 164 | 20 | 16 |
| % reads of most abundant OTU | 6.55 | 57.78 | 37.02 | 45.01 | 64.34 | 31.72 |

The fungal communities in the forest soil pyrocosms showed a fire response that is consistent with stimulation of a single genus of pyrophilous fungi, *Pyronema*, a reduction in richness and evenness of the community, but little apparent change to underlying composition (Table 2). Non-metric multidimensional scaling analysis shows that the two pyrocosm communities are about as distinct from each other as they are from the unburned control and also shows much variation occurs between individual depth samples and time points (Fig S2). However by sorting the OTU table by taxa that increase in the pyrocosms relative to the fire (File S2), three OTUs were found that exhibited dramatic increases in both pyrocosms relative to the control. All three were identified as *Pyronema* species (Table 2), a genus that is well known for its rapid fruiting following fire [4, 6].

**Table 2.**
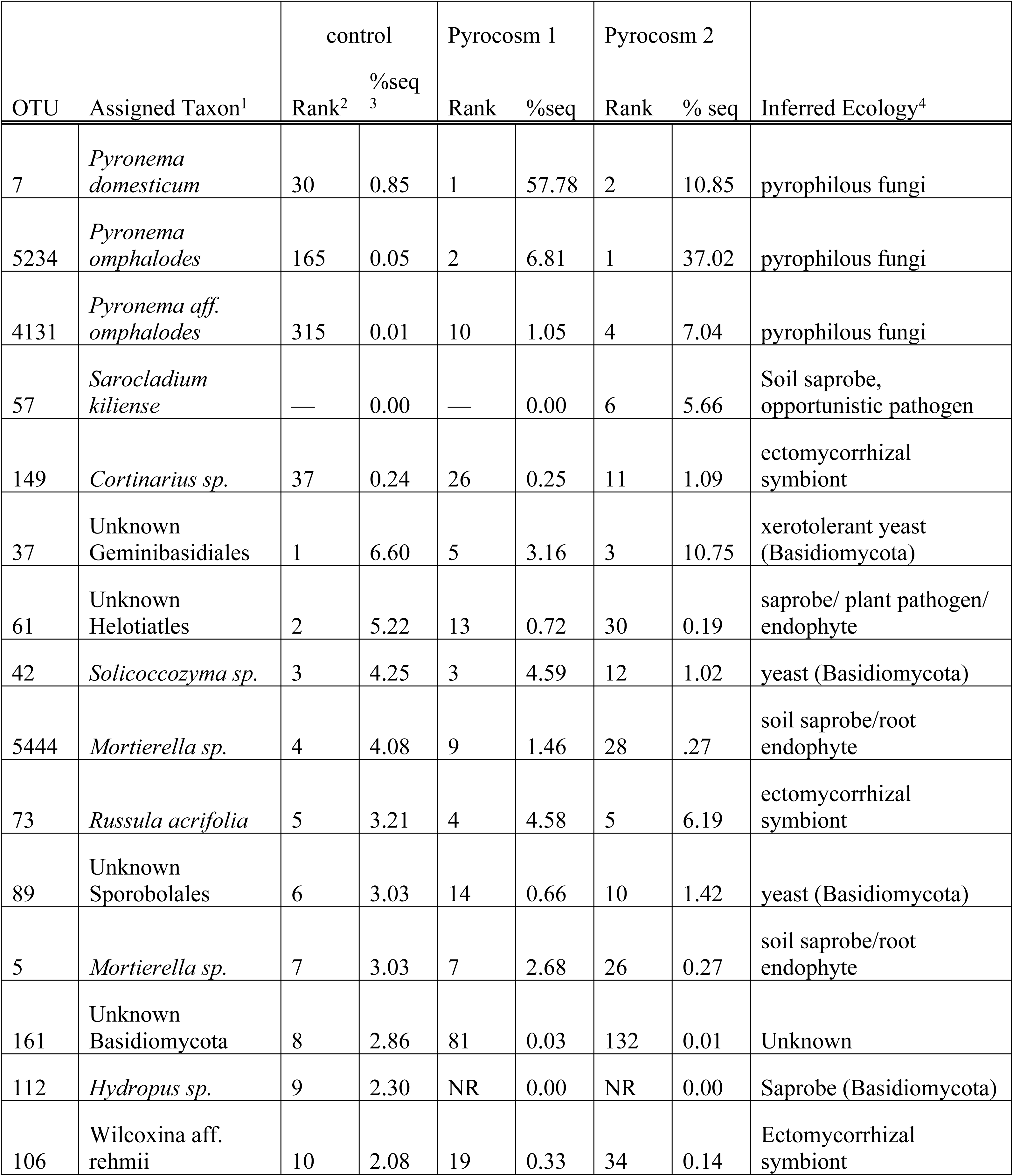
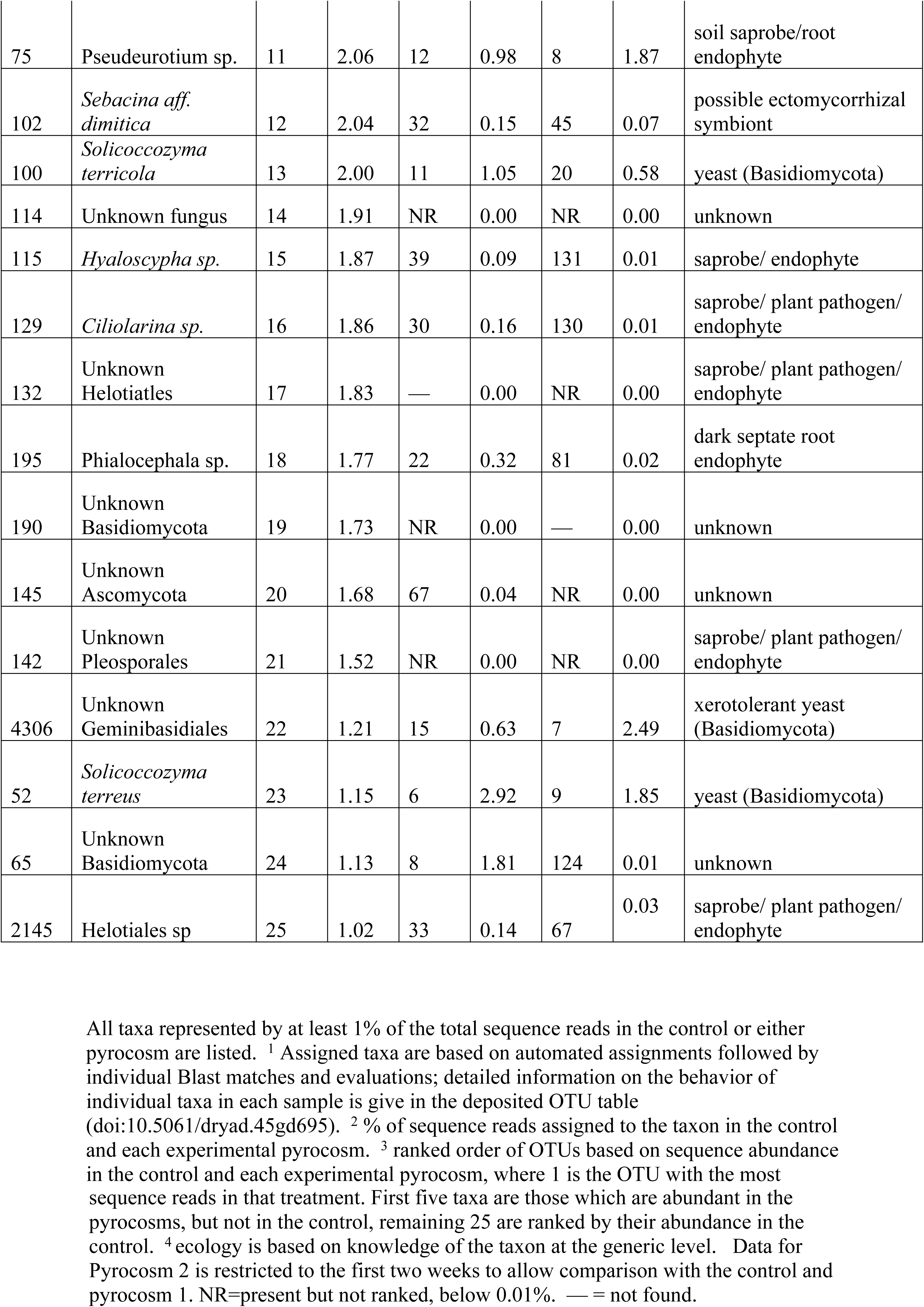
Most abundant OTUs summed across time and sample points.

| OTU | Assigned Taxon<sup>1</sup> | control |  | Pyrocasm 1 |  | Pyrocasm 2 |  | Inferred Ecology<sup>4</sup> |
| --- | --- | --- | --- | --- | --- | --- | --- | --- |
|  |  | Rank<sup>2</sup> | %seq<sup>3</sup> | Rank | %seq | Rank | % seq |  |
| 7 | *Pyronema domesticum* | 30 | 0.85 | 1 | 57.78 | 2 | 10.85 | pyrophilous fungi |
| 5234 | *Pyronema omphalodes* | 165 | 0.05 | 2 | 6.81 | 1 | 37.02 | pyrophilous fungi |
| 4131 | *Pyronema aff. omphalodes* | 315 | 0.01 | 10 | 1.05 | 4 | 7.04 | pyrophilous fungi |
| 57 | *Sarocladium kiliense* | — | 0.00 | — | 0.00 | 6 | 5.66 | Soil saprobe, opportunistic pathogen |
| 149 | *Cortinarius sp.* | 37 | 0.24 | 26 | 0.25 | 11 | 1.09 | ectomycorrhizal symbiont |
| 37 | Unknown Geminibasidiales | 1 | 6.60 | 5 | 3.16 | 3 | 10.75 | xerotolerant yeast (Basidiomycota) |
| 61 | Unknown Helotiatles | 2 | 5.22 | 13 | 0.72 | 30 | 0.19 | saprobe/ plant pathogen/ endophyte |
| 42 | *Solicoccozyma sp.* | 3 | 4.25 | 3 | 4.59 | 12 | 1.02 | yeast (Basidiomycota) |
| 5444 | *Mortierella sp.* | 4 | 4.08 | 9 | 1.46 | 28 | .27 | soil saprobe/root endophyte |
| 73 | *Russula acrifolia* | 5 | 3.21 | 4 | 4.58 | 5 | 6.19 | ectomycorrhizal symbiont |
| 89 | Unknown Sporobolales | 6 | 3.03 | 14 | 0.66 | 10 | 1.42 | yeast (Basidiomycota) |
| 5 | *Mortierella sp.* | 7 | 3.03 | 7 | 2.68 | 26 | 0.27 | soil saprobe/root endophyte |
| 161 | Unknown Basidiomycota | 8 | 2.86 | 81 | 0.03 | 132 | 0.01 | Unknown |
| 112 | *Hydropus sp.* | 9 | 2.30 | NR | 0.00 | NR | 0.00 | Saprobe (Basidiomycota) |
| 106 | *Wilcoxina aff. rehmii* | 10 | 2.08 | 19 | 0.33 | 34 | 0.14 | Ectomycorrhizal symbiont |
| 75 | *Pseudeurotium* sp. | 11 | 2.06 | 12 | 0.98 | 8 | 1.87 | soil saprobe/root endophyte |
| 102 | *Sebacina* aff. *dimitica* | 12 | 2.04 | 32 | 0.15 | 45 | 0.07 | possible ectomycorrhizal symbiont |
| 100 | *Solicoccozyma terricola* | 13 | 2.00 | 11 | 1.05 | 20 | 0.58 | yeast (Basidiomycota) |
| 114 | Unknown fungus | 14 | 1.91 | NR | 0.00 | NR | 0.00 | unknown |
| 115 | *Hyaloscypha* sp. | 15 | 1.87 | 39 | 0.09 | 131 | 0.01 | saprobe/ endophyte |
| 129 | *Ciliolarina* sp. | 16 | 1.86 | 30 | 0.16 | 130 | 0.01 | saprobe/ plant pathogen/ endophyte |
| 132 | Unknown Helotiales | 17 | 1.83 | — | 0.00 | NR | 0.00 | saprobe/ plant pathogen/ endophyte |
| 195 | *Phialocephala* sp. | 18 | 1.77 | 22 | 0.32 | 81 | 0.02 | dark septate root endophyte |
| 190 | Unknown Basidiomycota | 19 | 1.73 | NR | 0.00 | — | 0.00 | unknown |
| 145 | Unknown Ascomycota | 20 | 1.68 | 67 | 0.04 | NR | 0.00 | unknown |
| 142 | Unknown Pleosporales | 21 | 1.52 | NR | 0.00 | NR | 0.00 | saprobe/ plant pathogen/ endophyte |
| 4306 | Unknown Geminibasidiales | 22 | 1.21 | 15 | 0.63 | 7 | 2.49 | xerotolerant yeast (Basidiomycota) |
| 52 | *Solicoccozyma terreus* | 23 | 1.15 | 6 | 2.92 | 9 | 1.85 | yeast (Basidiomycota) |
| 65 | Unknown Basidiomycota | 24 | 1.13 | 8 | 1.81 | 124 | 0.01 | unknown |
| 2145 | Helotiales sp | 25 | 1.02 | 33 | 0.14 | 67 | 0.03 | saprobe/ plant pathogen/ endophyte |

All taxa represented by at least 1% of the total sequence reads in the control or either pyrocosm are listed. <sup>1</sup> Assigned taxa are based on automated assignments followed by individual Blast matches and evaluations; detailed information on the behavior of individual taxa in each sample is give in the deposited OTU table (doi:10.5061/dryad.45gd695). <sup>2</sup> % of sequence reads assigned to the taxon in the control and each experimental pyrocosm. <sup>3</sup> ranked order of OTUs based on sequence abundance in the control and each experimental pyrocosm, where 1 is the OTU with the most
sequence reads in that treatment. First five taxa are those which are abundant in the pyrocosms, but not in the control, remaining 25 are ranked by their abundance in the control. <sup>4</sup> ecology is based on knowledge of the taxon at the generic level. Data for Pyrocosm 2 is restricted to the first two weeks to allow comparison with the control and pyrocosm 1. NR=present but not ranked, below 0.01%. — = not found.

The three *Pyronema* OTUs were highly abundant in the burned soils, but not the controls (Table 2). Within two weeks of the fire, *Pyronema domesticum* (OTU7), was the most abundant taxon in pyrocosm 1, accounting for 67.78% of the sequence, and was the second most abundant sequence in pyrocosm 2, accounting for 10.85% of the sequence. *Pyronema omphalodes* (OTU 5234) was the second most dominant taxon in pyrocosm 1, accounting 6.81% of the total sequence, and it was the most dominant taxon in pyrocosm 2 in which it accounted for 37.02% of the aggregate sequence in the first two weeks. A third *Pyronema* (OTU 4131) ranked10^th^ the 4^th^ in abundance in the pyrocosms 1 and 2, respectively. All three *Pyronema* OTUs showed rapid rises, and declines in relative sequence abundance within the short time span of the experiment (Fig 3a). In addition, *Pyronema domesticum* (OTU7) fruited on the surface of the soil after 17 days in pyrocosm 2, where it was less abundant than *Pyronema omphalodes* (OTU 5234) (Fig 3b,c). All three *Pyromena* OTUs were detected in the control, but their read abundances were orders of magnitude lower and they did not increase with time as was the case in the pyrocosms (Fig 3). *Pyronema domesticum* (OTU7) was the most abundant of the three in the unburned control, where it accounted for 1.69% in the first week but declined to 0.01% by week two. As discussed below it also appeared in the mock community control presumably via tag switching [29] and accounted for 0.22% (92 reads) of the sequences.

**Fig 3.**
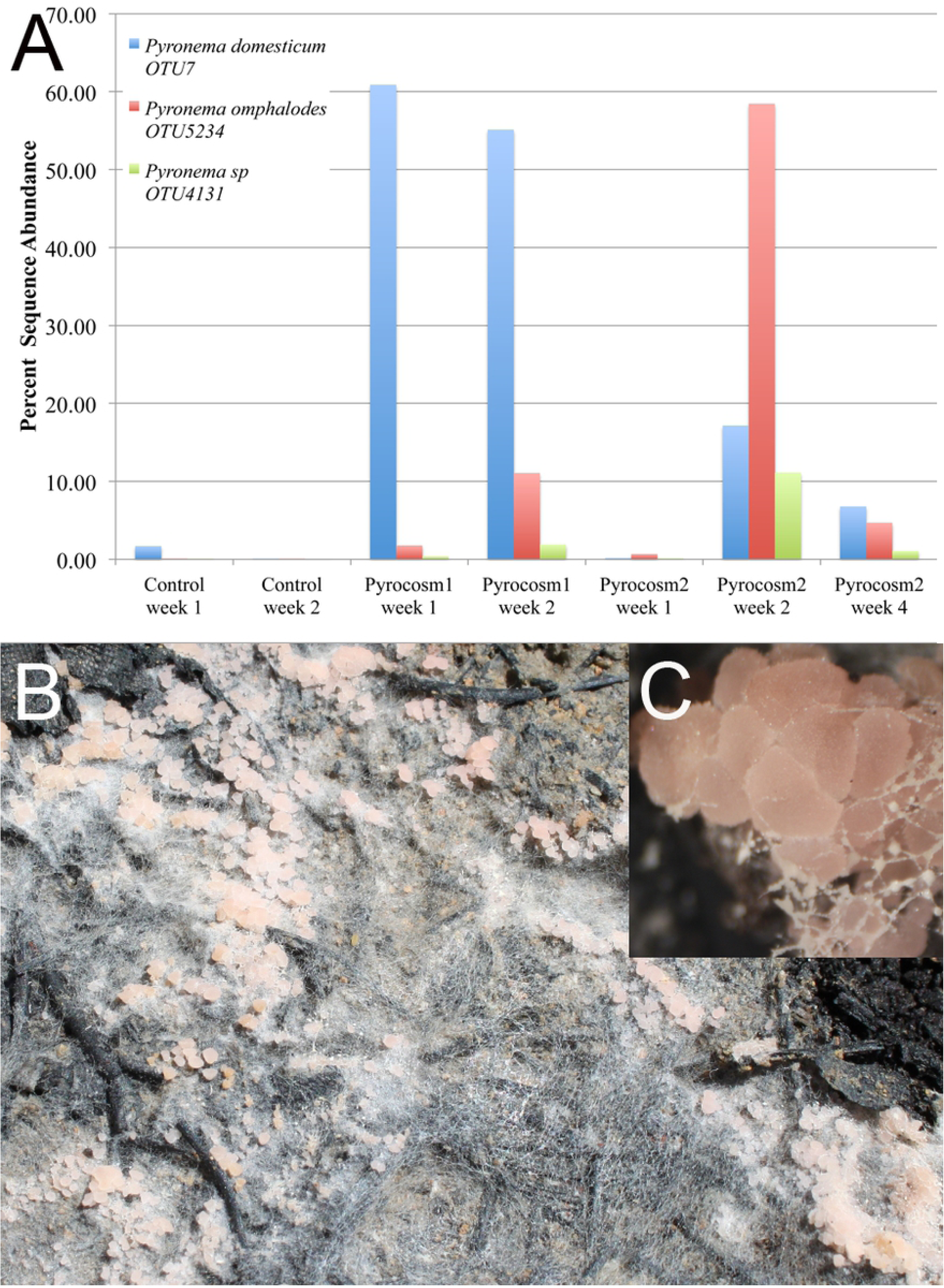
*Pyronema* species react rapidly to simulated fire in pyrocosms. A) post-fire percent sequence read abundance of three *Pyronema* OTUs is shown in the control, and pyrocosm 1 in weeks 1 and 2 and in pyrocosm 2 in weeks 1,2, and 4; B) mycelium and apothecia (ascocarps) of *Pyromena domesticum* fruiting on the surface of pyrocosm 2 just 17 days after the fire; C) closeup of same.

Only 79 OTUs had sequence abundance higher than 0.1% in at least one of the two fire pyrocosms. Of these, 38 OTUs showed nominal increases in sequence abundance relative to the unburned control in at least one of the pyrocosms, but only 10 OTUs showed increases in both pyrocosms relative to the control (File S2, column BR). Most of these increases were quite small, and only four OTUs increased by more than 1% relative abundance in both pyrocosms. Three of these were the *Pyronema* OTUs just discussed, the fourth was a *Russula* species (OTU73), an ectomycorrhizal fungus, that increased in sequence abundance by 1.37% and 3.7% in pyrocosms 1 and 2, respectively. The remaining six OTUs that increased in both pyrocosms, did so at much lower levels, and included two other ectomycorrhizal fungi (*Rhizopogon arctostaphyli* - OTU175, and an unknown *Thelephoraceae*-OTU879), two unidentified fungi (OTUs 304, 420), a Basidiomycota yeast (OTU631), and one additional *Pyronema* (OTU3739).

Other than *Pyronema*, differences in OTUs between the three units were subtle, often limited to a single experimental unit, and based primarily on differences in abundance of shared species, or the presence or absence of rarer taxa. For example, the top eight OTUs based on sequence abundance in the control were also found in both burned pyrocosms at similar levels, and of the top 25 most abundant OTUs in the control, all were detected in at least one of the two pyrocosms, although five of these were below the 0.01% threshold used for ranking (Table 2). One taxon, identified as *Sarocladium kiliense* (OTU 57, Hypocreales), accounted for 13.19% of the sequence in pyrocosm 2 in the first two weeks and was the 10^th^ most abundant taxon in that pyrocosm. It continued to increase and reached 45% of the sequence in the week four sample of pyrocosm 2, but it was absent from the control and pyrocosm 1.

Evenness and richness were reduced in the burned pyrocosms relative to the control (Fig 4C), but composition differences, other than those related to *Pyronema*, were not striking (table 2). The ranked abundance curves for the two pyrocosms are very similar, and distinct from that of the control (Fig 4). This pattern can also be seen in the level of dominance among component species. In unburned control the most abundant taxon, OTU 37 a xerotolerant yeast, represented 6.60% of the total sequence reads, and 25 OTUs each had more than 1% of the total reads. In contrast pyrocosm 1 and 2 had only 11 and 12 OTUs, respectively, that each had more than 1% of the reads. Among the less abundant taxa, the control pyrocosm had 394 OTUs that were each represented by at least 0.01% of the sequence, versus 158 and 155 OTUs in pyrocosms 1 and 2 respectively (Table 1, Fig 4).

**Fig 4.**
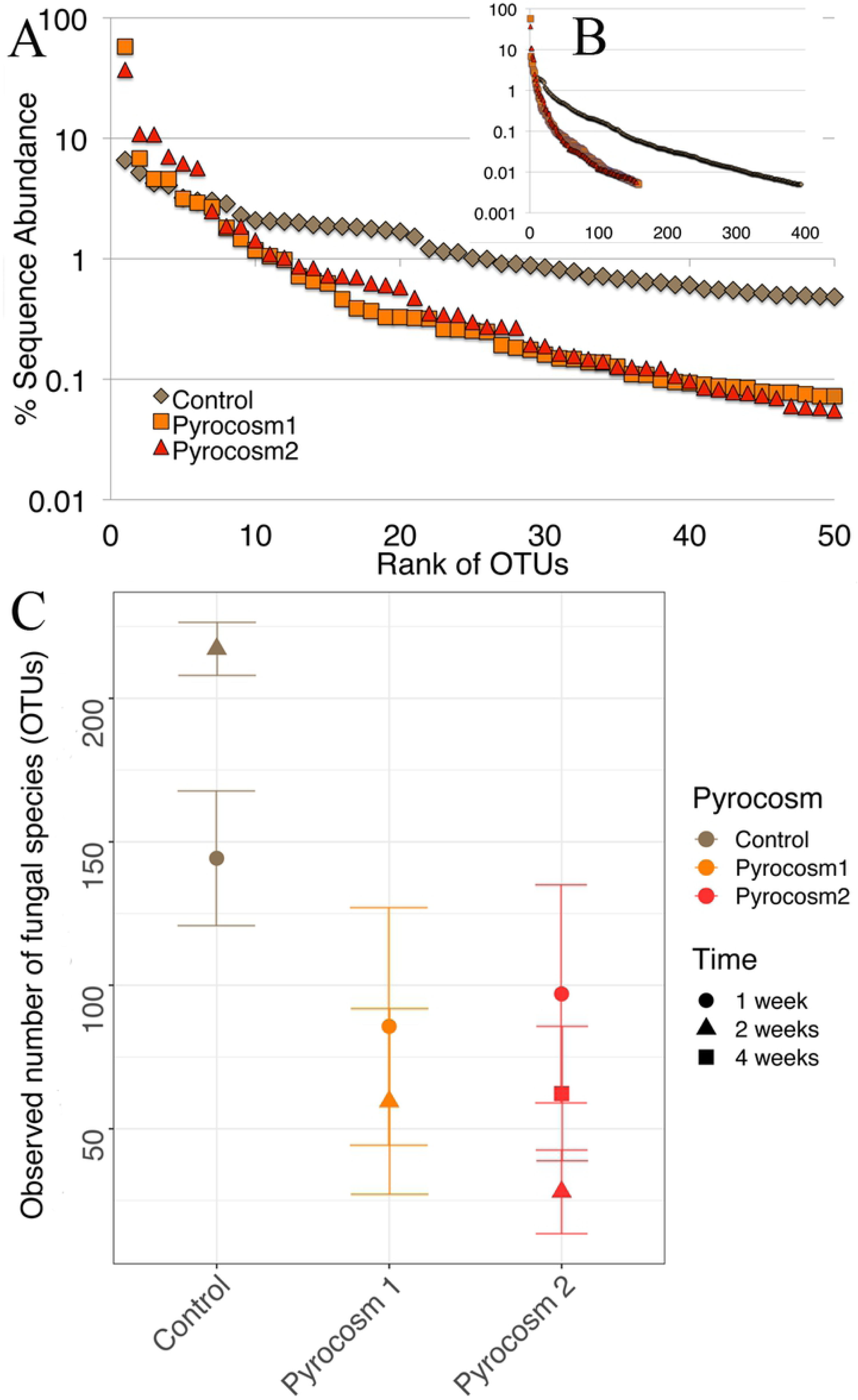
Differences in diversity of fungal communities within pyrocosms and control. A) Ranked abundance curves of pyrocosms and control based on relative read depth of for top 50 most abundant OTUs. B) Ranked abundance for all OTUs that are at least 0.01% of sequence. C) Richness estimates based on observed number of OTUs after rarifying samples to 3950 sequences; points show means for all samples; bars show SE

The four no DNA controls revealed low levels of contamination as is typical of high throughput sequence studies. Of the 10 OTUs that were based on more than 5 sequences in these controls, 6 were identified as *Morteriella sp.*(Mucoromycota), four were identified as an unknown Ascomycota in the Dothideomycetes or Sordariomycetes, one was a *Cortinarius* sp. (Basidiomycota), and one was an unidentified fungus. Three of the *Morteriella* OTUs (64, 108, 794), and the unidentified fungal OTU (544), were found at similar or greater levels in the no DNA controls compared to the experiment. Of the remaining contaminating OTUs, three occurred at much lower levels compared to the experimental samples (OTUs 1,5,14) and three were found in the experimental samples at levels below the thresholds used for analyses (i.e., 0.01%, or 5 sequences) (OTUs 38,556,1021). None of these contaminants are relevant to the results or analyses discussed above.

All the 10 knowns were recovered from the mock community controls, however, they yielded 16 OTUs with greater than 5 sequences each, and 20 OTUs with greater than 0.01% read abundance. This represents a 1.6 or 2-fold OTU inflation for these respective thresholds. This inflation was primarily due to minor amplicon variants of the known fungal species added, but also included some tag switching from the experimental samples. The most obvious case of the latter involved *Pyronema domesticum* (OTU7), which was the most abundant sequence in the experimental samples (290,128 total reads), and it accounted for 0.22% (92 reads) in the mock community control, even though it was not included in the community. Although the DNAs of the mock community were added in approximately equal concentrations *Rhodotorula mucilaginosa* (OTU13, Basidiomycota, yeast) accounted for 64.34% (27132 reads), and so its minor occurrence in the experimental samples (8 reads total), may be tag switching from control to experiment, although it is possible that it legitimately occurred in the samples.

## Discussion

We have shown that these very simple pyrocosms provide an experimental system in which soil heating can be achieved in predictable ways by simply altering fuel levels. Based on prior soil heating models [14–16], we would expect soils that differ in heat capacities and water contents to have different slopes (i.e. Fig 2b). However, the predictability within a given soil type should remain high, and we would expect the lognormal relationship between temperature and depth to remain. The strong correlation of peak temperature with log of depth, means that one can accurately predict temperatures at any depth if two points are measured, although the edge effects will broaden the range of temperatures achieved within a give depth zone. Earlier models have shown that water content of the soil will have the largest effect, particularly as it rises above 8% (Campbell et al 1994). This was specifically avoided in our studies because we assume that most large wildfires are likely to occur during very dry periods. We have also found that dry soil performed quite similarly to our sand pyrocosms, which helps with predicting peak temperatures in novel soils. However, if one were to model soil conditions of prescribed burns, higher water content of soils should be considered.

These results, and those of earlier modeling studies, have revealed features of the physical environment that are likely to be important for survival of propagules in the soil. The high heat capacity of soil increases with temperature [14], and means that soil heats slowly but retains heat for a long time (Fig 2a). This is shown by the fact that peak temperatures at depth in the pyrocosms are achieved hours after the fire has gone out. Essentially the heated surface soils have stored heat and become the source for heating deeper soils. This means that at depths below a few centimeters, soil organisms would be “slow-cooked”, with temperatures hovering around the peak for tens of minutes. This is important because the percent germination of spores of *Pyronema domesticum* and some other pyrophilous decrease as heat treatments persist for several minutes [7].

In our pyrocosms the large fuels (e.g. charcoal) transfer more heat to the surface and ultimately to the deeper soils, and were the best predictors of peak temperatures (Fig 2c). To relate this to a forest fire setting we envision a pattern of spotty heating across the landscape driven by the uneven distribution of coarse fuels. In highly intense crown fires, radiant heating of the surface soils are likely to contribute to much more soil heating than the litter [30, 31] and would likely even out some of this patchiness. Nevertheless, the uneven distribution of larger course fuels is likely to contribute to heating heterogeneity. The less homogeneous nature of natural soil compared to the sieved, dried, uniformity of the pyrocosm soil would add an additional source of spatial variation in peak soil temperatures [32].

To visualize this heating pattern and relate it back to soil organisms, let us couple this idea about uneven heating with a hypothesized ‘Goldilocks Zone’ – the not-too-hot, not-too-cool depths where the temperature range kills most organisms but is tolerated by propagules of pyrophilous species. For the sake of argument we will assume the Goldilocks zone exists between 50 and 70 °C. If we then couple the log relationship between soil depth and peak temperature, with the log relationship between mass of charcoal and peak temperature we can model this zone (Fig 5). This shows that the depth of the zone continues to increase with fuel load, while the thickness of the zone increases rapidly with addition of charcoal up to a threshold, but it then decreases slowly maintaining a similar width (∼3-5 cm, Fig 5b) over most of the modeled range. Note that the actual temperatures that define the zone might be hotter or cooler than assumed here and would likely vary with specific organisms [33], but different defining temperatures would only cause the zone to move up or down in depth and modify the width. However, the basic pattern of a thin selective zone that varies with local fuel load would still be valid. Therefore this model predicts that in a forest setting there would almost always be a survivable zone for pyrophilous microbes as long as the soil is deep enough.

**Fig 5.**
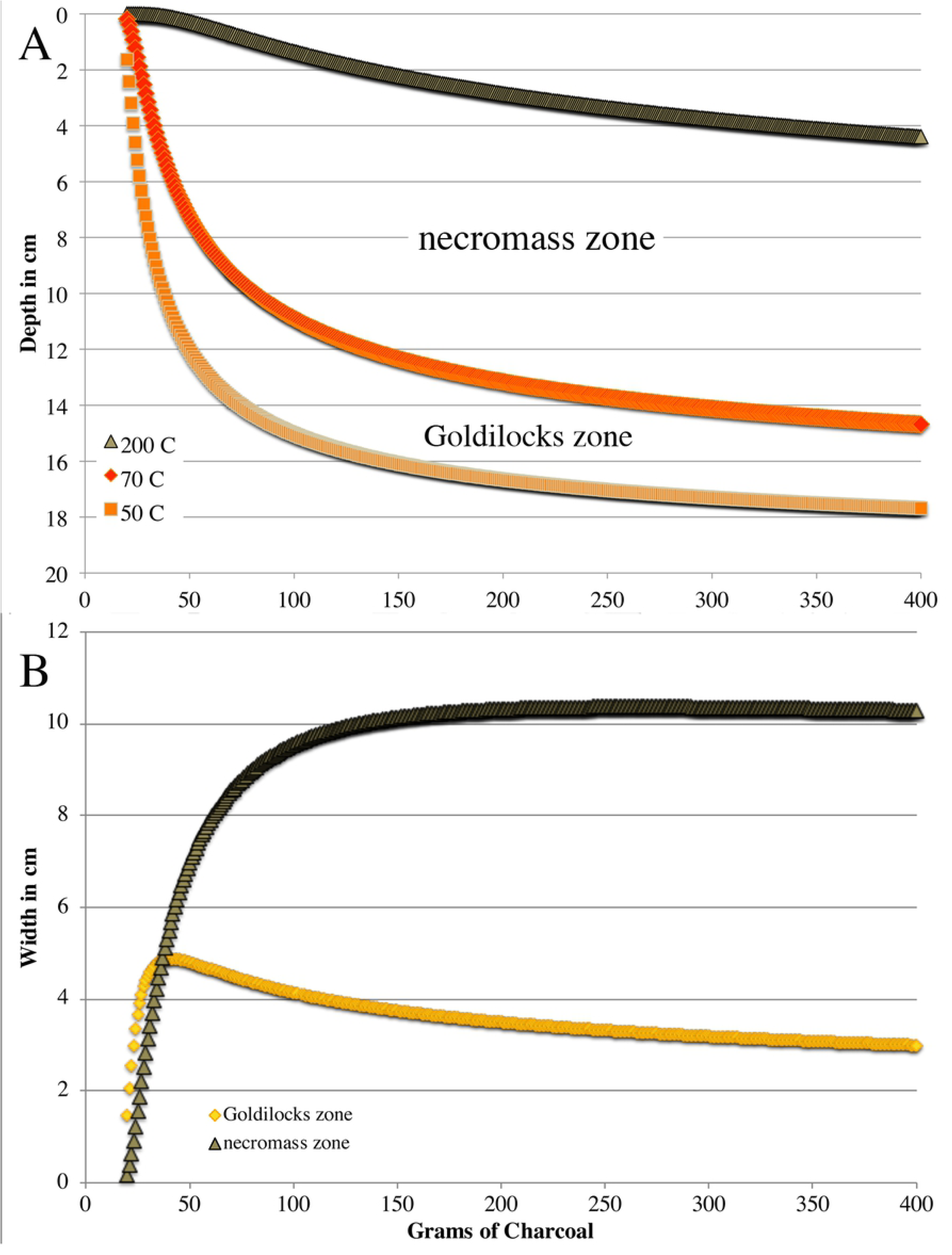
Modeled behavior of the “Goldilocks” and necromass zones as mass of charcoal is increased. **A)** depth at which peak temperatures reach 50, 70, and 200 °C are modeled by using the ln relationship between depth of peak temperature (Fig 1b), and the ln relationship between peak temperature at a given depth and mass of charcoal. The Goldilock and necromass zones are labeled between the hypothetical temperatures that define them B) Width of both zones over the same range of charcoal mass.

Post-fire soil chemistry is correlated with temperature and therefore creates additional depth-structured zones that recolonizing organisms will encounter. Temperatures greater than 350 °C cause heavy pyrolyzation of cellulose and lignin resulting in highly aromatic forms of biochar [17] that are relatively recalcitrant to microbial degradation [34]. According to our regression-based model these temperatures would only occur within about a centimeter of the surface in the highest fuel loads tested, and would be non-existent in the lower fuel loads tested (File S3). At soil temperatures between 200 and 300 °C pyrolyzation begins, but is more limited in the extent of aromatic conversion [17]. Figure 5A shows the gradual increase in depth of 200 °C peak temperature as course fuel is increased. Soil temperatures in higher than 175 to 200 °C volatilize waxes and lipids that then condense at lower temperatures in the soil below and cause a hydrophobic layer that can facilitate erosion [35]. It is not known if these concentrated hydrophobic materials serve as energy-rich carbon sources for microbes, but it is known that disappear with time [35], and we think it would be surprising if microbes were not involved in some way. Temperatures lower than this but above the lethal temperature produce only modest chemical changes, but effectively kill most organisms. We define this as a necromass zone, where death of soil flora, fauna, and microbes results in the release of labile nutrients with minimal pyrolyzation. This should be a rich habitat for pyrophilous organisms that survive in the slightly cooler Goldilocks zone below it and then grow rapidly into the necromass rich habitat after fire. If we model the depth and width of this zone in relation to coarse fuel, we see that the depth of the necromass zone drops gradually (Fig 5A), while the width of the zone increases rapidly up to a threshold (∼100 g of charcoal, about 4 briquettes), and then stays relatively constant (Fig 5B). Here we have defined the necromass zone as the area between peak temperatures of 70 and 200 °C, but even if one defined the bordering temperatures differently we would expect the basic trends to remain.

The rapid, massive response of *Pyronema* species in both fire pyrocosms (Fig 3) demonstrates that this system is capable of simulating at least some known pyrophilous microbes. *Pyronema*’s response in our system is impressive in that three OTUs accounted for 55 to 65% of all fungal sequence reads in both pyrocosms within just two weeks of the fire, and the most dominant *Pyromena* OTU in each pyrocosm accounted for 37 to 58 % of all reads in the two-week post-fire period, compared to less than 1% in the control (Table 2). For comparison the most dominant fungus in the unburned control was a xerotolerant yeast that accounted for less than 7% of the sequence (Table 2). This latter value is typical of dominants in other studies of soil fungi in which the most abundant species typically account for only few percent of the sequences, and all species with more than 0.1% are viewed as dominants [36]. *Pyronema* species achieved a dominance that is roughly 6 to 10-fold higher than is typical of the most dominant fungal soil fungi, and they did so in just one or two weeks.

*Pyromena* species have been known to be rapid responders to fire for over a century [4], but all of the earlier literature is based on the fruiting response of the species. Evidence of its *in situ* mycelial response was lacking. *In vitro* studies have shown that *Pyronema* species grow rapidly in situations where there are few competitors [7, 37]. *Pyronema* was also mentioned in Petersen’s (1970) experimental fire work, but its fruiting, though rapid, was not consistent enough for it to be considered in the successional patterns that Petersen discussed. Interestingly in our study, it only fruited in one of the two pyrocosms, even though it dominated in both. Thus, it may be much more common in post-fire soils then its fruiting record suggests.

At least some pyrophilous fungi appear to have dormant spores or sclerotia in the soil, and these germinate following heating or chemical changes associated with fire [6, 38]. This mode of activating quiescent propagules is consistent with the rapid response we saw, but we did not specifically test for it by preventing dispersal. Nevertheless, the presence of all three dominant *Pyronema* OTUs in both pyrocosms shows that no inoculation was necessary for a rapid response.

The small response of *Pyromena domesticum* in week one of the control is interesting (Fig 3), and corresponded to 1.69% (3535 total reads) of the sequence in that sample. This is three orders of magnitude higher that the 92 sequence reads that contaminated the mock community, and therefore must be a real response of *Pyronema* within the unburned control soil. We interpret this as a short-term simulation following the wetting of the dried soil. One could imagine that such transient responses could take advantage of brief periods of low competition to renew soil inoculum between fire events. In any case, the prevalence of *Pyromena domesticum* in the control dropped to 0.006% by week two. The burned pyrocosms also show rapid increases and decreases in *Pyronema* within one-week periods (Fig 3). This rapid turnover shows that most of the *Pyronema* DNA in the soil does not linger long, and it suggests either autolysis or degradation by other components of the microbial community. In contrast the presence of ectomycorrhizal taxa such as *Cortinarius* and *Russula spp.* that occur in similar levels within the control and burned pyrocosms (Table 2) are likely due to survival of environmental DNA, perhaps as spores, as neither taxon would be expected to grow without a host.

It is less clear if fungi other than *Pyronema* specifically responded to the experimental fire in a positive way. The inclusion of ectomycorrhizal fungi in the set of fungi that appeared to increase after the fire at low levels (Table 2) shows that these apparent changes are within the level of stochastic change, as growth of these ectomycorrhizal fungi could not have occurred without a host tree. For most other fungi we have no knowledge of their autecologies, so when they appear to respond at similar levels to the ectomycorrhizal fungi there is no strong evidence for or against their fire response. Longer incubations times and increased replication are necessary to resolve this. For example, the continued increase of *Sarocladium kiliense* (OTU 57) up to 45% of the sequence at the four week sampling time, shows that this fungus did grow following the fire treatment, even though our limited knowledge of its ecology does not give us reason to expect a post-fire response. However, its presence in only one of the two pyrocosms means that we cannot draw generalizations about its response without substantially increased replication.

The negative effects of fire on most fungi is clear from the reduced species richness and shape of the ranked abundance curves (Fig 4). This result is concordant with a meta-analysis of fungal response to fire [3], and makes intuitive sense based on the killing effect of soil heating. However, having taxa with extremely high read abundance can distort perception of community structure because our ability to detect less abundant taxa is reduced as more of sequence depth is consumed by the dominants [39]. The high abundance of *Pyronema* in these experiments, certainly falls within this realm, and the high OTU overlap between the control and the burned pyrocosms (Table 2) makes the possibility of an artifactual reduction in richness plausible. However, if we remove the sequence reads from the top three *Pyronema* OTUs, and recalculate percent dominance, the rank abundance curves from the burned pyrocosms are still much less even than that of the control (Fig S3). This shows that in addition to *Pyronema* other taxa are also driving the shape of the ranked abundance curves, and thus the reduction in richness and evenness are unlikely to be artifacts.

We propose that the pyrocosm system is an excellent model system to dissect the post-fire microbial community, and this can be done in a variety of ways. For example soil source, fire intensity, watering regime, access to external inoculum and incubation conditions can all be varied and replicated to study the process of post-fire community assembly. If pyrocosm experiments were incubated longer or under different conditions (e.g., water, soil type, temperature), other common, known pyrophilous fungi [6] might develop from the natural inoculum just as *Pyronema* did. However, even if they did not, most of them grow well in culture [7] and could be added into post-fire soils that lack them in varying orders and combinations to determine environmental versus biological interactions that underlie community assembly [40]. There are also indications of parallel postfire responses in bacteria and microfauna [2] that could be studied in similar ways with this system.

Finally, we propose that *Pyronema* is an excellent model organism for the study of fire fungal ecology. In addition to their ease of isolation and rapid growth rates, three genomes from *Pyronema* species have now been completely sequenced, assembled and annotated, and are available on Mycocosm (http://jgi.doe.gov/fungi), the genome portal for the U.S. Department of Energy (DOE) Joint Genome Institute (JGI) [41]. One of these *Pyronema* species had been sequenced and assembled previously [42], and two are new. Coupling these genomic resources with the rapid dominance of *Pyronema* in postfire soil can thus be used to explore the functional roles of this fungus in a realistic environment. For example, gene expression could be studied by incubating *Pyronema* on temperature defined soil layers removed from pyrocosms (File S2) followed by extraction and sequencing of mRNA. This approach would help improve our understanding of what these fungi live on within the post-fire soil.

Furthermore, *Pyronema* is not unique within the post-fire community; genomes of 10 other pyrophilous fungi have now been sequenced, assembled, and annotated (Table 3). The list includes representatives of most of the common genera of pyrophilous fungi [6]. This is important because metatranscriptomic approaches with fungi are generally limited by the number and taxonomic coverage of sequenced and annotated genomes [43]. Due to its relative simplicity, the post-fire soil fungal community may now have the best genomic resources of any soil system, and these resources can be used in combination with the pyrocosms to dissect interacts with post-fire soil chemistry or with other organisms in this environment.

**Table 3.**
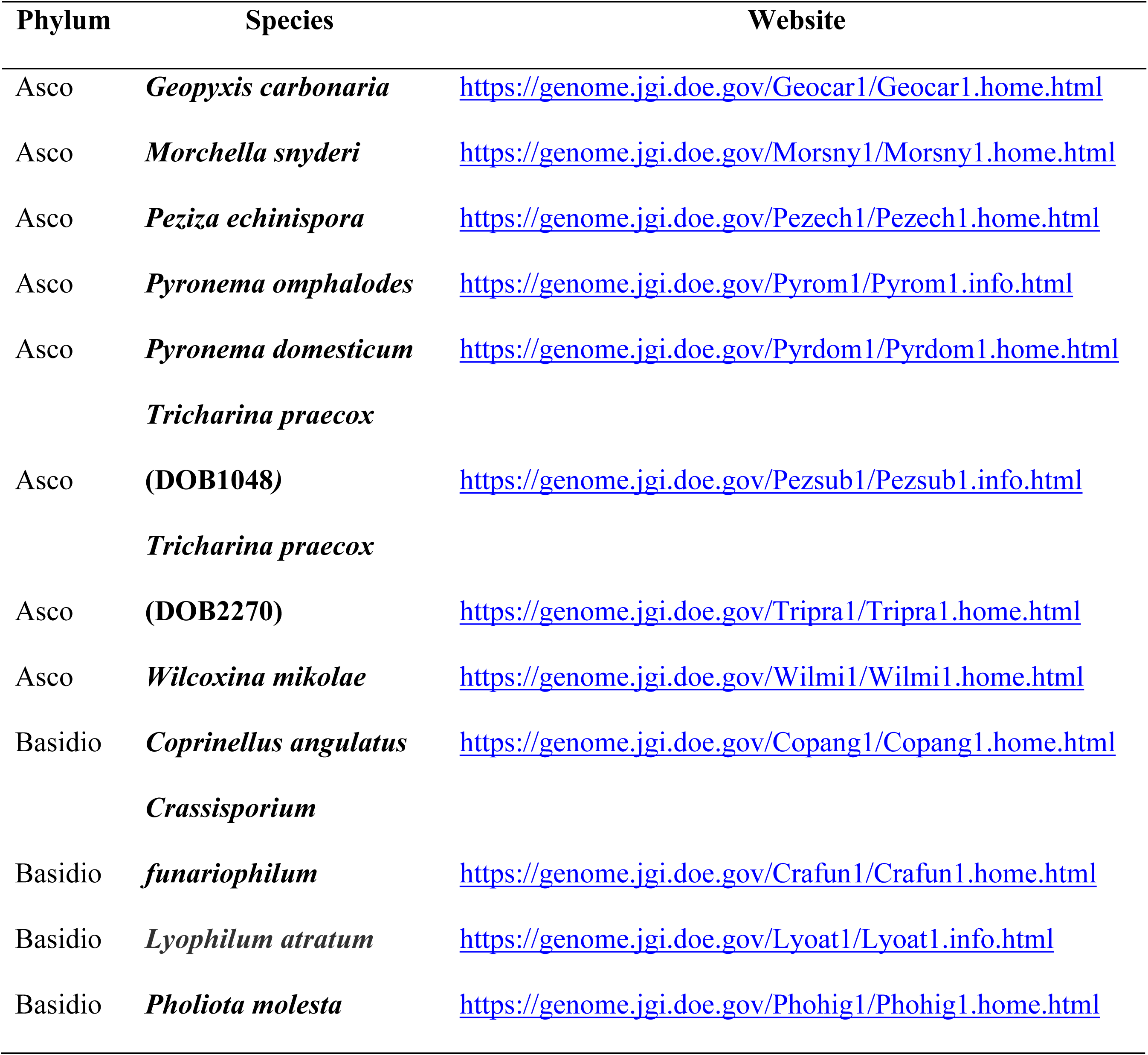
Sequenced, Assembled and Annotated Genomes of Pyrophilous Fungi

## Conclusions

Pyrocosms now add to the growing list of experimental systems that can be used to study fungal community ecology [44], and they specifically relate to a natural environment that is growing in importance as fire size and intensity continue to rise [45]. Furthermore the community and the factors affecting this environment are simple enough that experimental and modeling approaches might be easily combined to achieve greater predictability [46], while the availability of genomic resources open avenues to dissect the functional roles of the component species in great detail.

## Acknowledgments

Funding was provided by Department of Energy grant DE-SC0016365 to TDB, and genome sequencing was accomplished through the a community sequencing project from the Joint Genome Institute to TDB.

## Supporting information

**Fig S1. Effect of flash fuels is subtle.** 172 gm of flash fuels caused temperatures to rise slightly at 10.5 cm below the surface, but ultimately achieved the same peak temperature as solar heating in an unburned pyrocosm monitored simultaneously. Pyrocosms with the same flash fuel load, but one or two charcoal briquettes (25-51 g) heated more rapidly and achieve higher peak temperatures.

**Fig S2. NMDS plots using Bray-Curtis dissimilarity**. Samples were rarified to 3950 reads/sample. Adonis R^2^ = 0.33 and 0.19 for Bray-Curtis (A) or Jaccard (B) respectively.

**Fig S3. Ranked abundance curves with three *Pyronema* OTUs dropped and percent read abundance recalculated without them.** Top 50 most abundant OTUs are shown.

**File S1 - Additional notes on pyrocosm assembly and use**

## Data Availability

Temperature data for all pyrocosms and the complete OTU table are deposited in Dryad: doi:10.5061/dryad.45gd695; Representative sequences for all the OTUs are deposited in NCBI XXXXXX-XXXXXX, and the raw read data are deposited in the short read archive: PRJNA559408.

## References

1. Keeley JE, Pausas JG, Rundel PW, Bond WJ, Bradstock RA. Fire as an evolutionary pressure shaping plant traits. Trends in Plant Science. 2011;16(8):406–11.

2. Pressler Y, Moore JC, Cotrufo MF. Belowground community responses to fire: meta-analysis reveals contrasting responses of soil microorganisms and mesofauna. Oikos. 2019;128(3):309–27.

3. Dove NC, Hart SC. Fire reduces fungal species richness and in situ mycorrhizal colonization: a meta-analysis Fire Ecology. 2017;13(2):37–65.

4. Seaver FT. Studies in Pyrophilous Fungi: I. The Occurrence and Cultivation of Pyronema. Mycologia. 1909;1(4):131–9.

5. Lisiewska M. Macrofungi on special substrates. In: Winterhoff W, editor. Fungi in vegetation science. Dordreckt, The Netherlands: Kluwer Academic Publishers; 1992. p. 151–82.

6. Petersen PM. Danish Fireplace Fungi. Dansk Botanisk Arkiv. 1970;27(3):1–97.

7. Elabyad MSH, Webster J. Studies on Pyrophilous Discomycetes 1. Comparative Physiological Studies. Transactions of the British Mycological Society. 1968;51:353-&.

8. Emerson M. Chemical Activation of Ascospore Germination in Neurospora crassa. Journal of Bacteriology. 1948;55:327–30.

9. Baar J, Horton TR, Kretzer AM, Bruns TD. Mycorrhizal colonization of Pinus muricata from resistant propagules after a stand-replacing wildfire. New Phytologist. 1999;143(2):409–18.

10. Glassman SI, Levine CR, DiRocco AM, Battles JJ, Bruns TD. Ectomycorrhizal fungal spore bank recovery after a severe forest fire: some like it hot. Isme Journal. 2016;10(5):1228–39.

11. Peay KG, Garbelotto M, Bruns TD. Spore heat resistance plays an important role in disturbance-mediated assemblage shift of ectomycorrhizal fungi colonizing Pinus muricata seedlings. Journal of Ecology. 2009;97(3):537–47.

12. Peay KG, Bruns TD, Garbelotto M. Testing the ecological stability of ectomycorrhizal symbiosis: effects of heat, ash and mycorrhizal colonization on Pinus muricata seedling performance. Plant and Soil. 2010;330(1-2):291–302.

13. Izzo A, Canright M, Bruns TD. The effects of heat treatments on ectomycorrhizal resistant propagules and their ability to colonize bioassay seedlings. Mycological Research. 2006;110:196–202.

14. Massman WJ. Modeling soil heating and moisture transport under extreme conditions: Forest fires and slash pile burns. Water Resources Research. 2012;48.

15. Massman WJ, Frank JM. Effect of a controlled burn on the thermophysical properties of a dry soil using a new model of soil heat flow and a new high temperature heat flux sensor. International Journal of Wildland Fire. 2004;13(4):427–42.

16. Campbell GS, Jungbauer JD, Bidlake WR, Hungerford RD. Prediction the effect of temperature on soil thermal-Conductivity Soil Science. 1994;158(5):307–13.

17. Keiluweit M, Nico PS, Johnson MG, Kleber M. Dynamic Molecular Structure of Plant Biomass-Derived Black Carbon (Biochar). Environmental Science & Technology. 2010;44(4):1247–53.

18. Certini G. Effects of fire on properties of forest soils: a review. Oecologia. 2005;143(1):1–10.

19. Garcia-Chevesich P, Pizarro R, Stropki CL, de Arellano PR, Ffolliott PF, DeBano LF, et al. Formation of Post-fire Water-Repellent Layers in Monterrey Pine (Pinus radiata D. DON) Plantations in South-Central Chile. Journal of Soil Science and Plant Nutrition. 2010;10(4):399–406.

20. Mainwaring K, Hallin IL, Douglas P, Doerr SH, Morley CP. The role of naturally occurring organic compounds in causing soil water repellency. European Journal of Soil Science. 2013;64(5):667–80.

21. Schoch CL, Seifert KA, Huhndorf S, Robert V, Spouge JL, Levesque CA, et al. Nuclear ribosomal internal transcribed spacer (ITS) region as a universal DNA barcode marker for Fungi. Proceedings of the National Academy of Sciences of the United States of America. 2012;109(16):6241–6.

22. Smith DP, Peay KG. Sequence Depth, Not PCR Replication, Improves Ecological Inference from Next Generation DNA Sequencing. Plos One. 2014;9(2).

23. Edgar RC. UPARSE: highly accurate OTU sequences from microbial amplicon reads. Nature Methods. 2013;10(10):996-+.

24. Caporaso JG, Kuczynski J, Stombaugh J, Bittinger K, Bushman FD, Costello EK, et al. QIIME allows analysis of high-throughput community sequencing data. Nature Methods. 2010;7(5):335–6.

25. Koljalg U, Nilsson RH, Abarenkov K, Tedersoo L, Taylor AFS, Bahram M, et al. Towards a unified paradigm for sequence-based identification of fungi. Molecular Ecology. 2013;22(21):5271–7.

26. Altschul SF, Gish W, Miller W, Myers EW, Lipman DJ. BASIC LOCAL ALIGNMENT SEARCH TOOL. Journal of Molecular Biology. 1990;215(3):403–10.

27. Kircher M, Sawyer S, Meyer M. Double indexing overcomes inaccuracies in multiplex sequencing on the Illumina platform. Nucleic Acids Research. 2012;40(1).

28. R_core_team. R: A language and environment for statistica computing. Vienna, Austria: R foundation for Statistical Computing; 2014.

29. Carlsen T, Aas AB, Lindner D, Vralstad T, Schumacher T, Kauserud H. Don’t make a mista(g)ke: is tag switching an overlooked source of error in amplicon pyrosequencing studies? Fungal Ecology. 2012;5(6):747–9.

30. Santin C, Doerr SH, Merino A, Bryant R, Loader NJ. Forest floor chemical transformations in a boreal forest fire and their correlations with temperature and heating duration. Geoderma. 2016;264:71–80.

31. Thompson DK, Wotton BM, Waddington JM. Estimating the heat transfer to an organic soil surface during crown fire. International Journal of Wildland Fire. 2015;24(1):120–9.

32. Stoof CR, De Kort A, Bishop TFA, Moore D, Wesseling JG, Ritsema CJ. How Rock Fragments and Moisture Affect Soil Temperatures during Fire. Soil Science Society of America Journal. 2011;75(3):1133–43.

33. Pingree MRA, Kobziar LN. The myth of the biological threshold: A review of biological responses to soil heating associated with wildland fire. Forest Ecology and Management. 2019;432:1022–9.

34. Singh N, Abiven S, Torn MS, Schmidt MWI. Fire-derived organic carbon in soil turns over on a centennial scale. Biogeosciences. 2012;9(8):2847–57.

35. DeBano LF. The role of fire and soil heating on water repellency in wildland environments: a review. Journal of Hydrology. 2000;231:195–206.

36. Egidi E, Delgado-Baquerizo M, Plett JM, Wang JT, Eldridge DJ, Bardgett RD, et al. A few Ascomycota taxa dominate soil fungal communities worldwide. Nature Communications. 2019;10.

37. Elabyad MSH, Webster J. Studies on Pyrophilous Discomycetes 2. Competition. Transactions of the British Mycological Society. 1968;51:369-&.

38. Jalaluddin M. Studies on Rhizina undulata I. mycelial grrowth and ascospore germination Transactions of the British Mycological Society. 1967;50:449-+.

39. Adams RI, Amend AS, Taylor JW, Bruns TD. A Unique Signal Distorts the Perception of Species Richness and Composition in High-Throughput Sequencing Surveys of Microbial Communities: a Case Study of Fungi in Indoor Dust. Microbial Ecology. 2013;66(4):735–41.

40. Kraft NJB, Adler PB, Godoy O, James EC, Fuller S, Levine JM. Community assembly, coexistence and the environmental filtering metaphor. Functional Ecology. 2015;29(5):592–9.

41. Nordberg H, Cantor M, Dusheyko S, Hua S, Poliakov A, Shabalov I, et al. The genome portal of the Department of Energy Joint Genome Institute: 2014 updates. Nucleic Acids Res. 2014;42:(1):D26–31..

42. Traeger S, Altegoer F, Freitag M, Gabaldon T, Kempken F, Kumar A, et al. The Genome and Development-Dependent Transcriptomes of Pyronema confluens: A Window into Fungal Evolution. Plos Genetics. 2013;9(9).

43. Kuske CR, Hesse CN, Challacombe JF, Cullen D, Herr JR, Mueller RC, et al. Prospects and challenges for fungal metatranscriptomics of complex communities. Fungal Ecology. 2015;14:133–7.

44. Bruns TD. The developing relationship between the study of fungal communities and community ecology theory. Fungal Ecology. 2019;39:393–402.

45. Westerling AL, Hidalgo HG, Cayan DR, Swetnam TW. Warming and earlier spring increase western US forest wildfire activity. Science. 2006;313(5789):940–3.

46. Widder S, Allen RJ, Pfeiffer T, Curtis TP, Wiuf C, Sloan WT, et al. Challenges in microbial ecology: building predictive understanding of community function and dynamics. Isme Journal. 2016;10(11):2557–68.

